# RhythmWriter: GPU-accelerated Contractility Quantification for Large Datasets

**DOI:** 10.64898/2026.10.15.759435

**Authors:** Brandon Perestrelo, Marie Lecocq, Alvin Shrier, Gil Bub

**Affiliations:** Department of Physiology, McGill University, Montreal, Quebec, Canada

**Keywords:** Cardiomyocyte Contractility Algorithm, GPU-Accelerated High-Throughput Analysis, Open-Source FIJI Plugin, hiPSC-CM Aggregates

## Abstract

Human induced pluripotent stem cell-derived cardiomyocytes (hiPSC-CMs) have been considered optimal model systems for use in cardiac safety testing, yet their adoption remains low, largely because of technical limitations in the analysis methodology. RhythmWriter (RW) is an open-source GPU plugin for ImageJ developed to quantify contractile motion over extended periods of time. RW compensates for baseline drift, is time-scale invariant and is capable of very rapid data analysis. As a consequence, RW is capable of generating simultaneous high-resolution contraction restitution curves from multiple aggregates of hiPSC-CMs derived from premature-stimulus (S1S2) and dynamic-ramp restitution protocols. The aggregates exhibit brief refractory periods, which allow the induction of S2 beats at coupling intervals down to 130 ms and 1:1 entrainment during periodic stimulation down to a cycle length of 245 ms. Simulations confirm the ability of RW to track programmed contractile strength and stimulation intervals even at rapid pacing rates. This makes possible, for the first time, high-throughput functional assessment of hiPSC-CM aggregates for contractility drug screening and maturation.

## Introduction

The Comprehensive in vitro Proarrhythmia Assay (CiPA), an international initiative launched by the US Food and Drug Administration (FDA) alongside industry and academic partners, set out to implement a framework to more accurately assess the risk of the potentially fatal cardiac arrhythmia torsade de pointes (TdP) [1–3]. Human induced pluripotent stem cell-derived cardiomyocytes (hiPSC-CMs) retain the genetic makeup of the donor [4, 5] which makes them a highly suitable in vitro cardiomyocyte model [6, 7]. A blinded multisite study found that hiPSC-CMs could be used to accurately predict TdP risk [8] and subsequently, the International Council for Harmonization (ICH) accepted hiPSC-CM assays as a permissible complement to using animal and immortalized cell lines for preclinical cardiac safety assessment [9, 10] and proarrhythmic risk [11]. This coincides with the FDA’s continued push to reduce unnecessary animal testing [12]. However, fewer than 1% of new drug applications to the FDA include any hiPSC-CM data [10]. This is due in part to technical difficulties generating and analyzing relevant electrical or mechanical data in a high throughput fashion [13, 14]. Manual patch clamp of hiPSC-CMs fails to meet the throughput required for drug development [7, 15, 16], while high throughput patch clamp has not evolved to a level sufficient to provide robust screening of hiPSC-CMs. Methods have been developed to quantify the motion of contracting cardiac tissue non-invasively, as a proxy for contractile force [17–22]. Since contraction lies downstream of electrical activity and calcium cycling [23], assessing motion has the potential to identify disease and drug phenotypes [21, 22].

Contractility assays developed to date that are technically feasible [17] can provide important insights into cardiomyocyte function. MUSCLEMOTION (MM), a popular ImageJ macro, was shown by machine-learning validation, to be able to distinguish drug phenotypes better than optical flow [24]. However, MM is best suited for short recording periods (*<*5 s) because for longer recording periods MM is limited due to computational speed and baseline drift [25–27]. Another more recent method, PLoRI, attempts to deal with baseline drift by assessing the rolling intensity displacement from the median [28]. However, at more rapid pacing rates the median tends to rise compromising the analysis. An alternate approach, used in a number of studies, is to assess the rate of change of displacement [17, 18, 20, 21], which is immune to drift and scalable. However, this approach fails to distinguish between the speed of contraction and relaxation [17, 20] and does not manage lengthening of contractile complexes [21].

Mechanical restitution, the increase in the force of contraction as a function of the recovery interval, was initially studied to gain insights about cardiac calcium cycling dynamics [29, 30] before intracellular calcium could be measured directly [31, 32]. Mechanical restitution, which employs optical techniques to assess contraction of isolated animal myocytes [33–36] and heart cell aggregates [37], has included two complementary protocols. The S1S2 protocol assesses restitution by introducing a stimulus of increasing prematurity (S2) following a steady-state train (S1) [29, 33]. The dynamic ramp protocol assesses steady state contractile force at increasing pacing frequencies changed in a stepwise fashion, thereby generating a force–frequency relationship [34, 38]. In hiPSC-CMs, restitution protocols have been applied to action potentials [39] and calcium transients [40]. While the relationship between frequency and force has been assessed in engineered tissues at a few rates [41, 42], mechanical restitution of hiPSC-CMs remains to be characterized.

All of the assays developed to date have limited applicability for drug screening because of their inability to simultaneously acquire multiple hiPSC-CM preparations and analyze the data at high sampling speeds over extended periods of time [18, 25]. RhythmWriter (RW) is an open-source plugin for Fiji (ImageJ) [43] that overcomes these limitations by enabling the analysis of multiple electrically paced hiPSC-CM aggregate preparations over sufficiently long-time intervals to allow characterization of contraction restitution properties. Here, we use RW to measure S1S2 and dynamic restitution of contraction in several hiPSC-CM aggregates paced in parallel for over 40 minutes.

## Results

### RhythmWriter Algorithm

Numerous methods quantify this motion as the mean pixel difference between each frame of a recording and a reference frame [21, 25]:

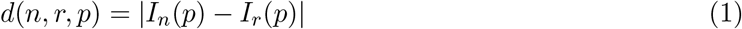

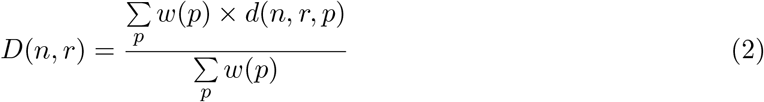

where *I*_*n*_(*p*) is the intensity of pixel *p* in frame *n, r* is the reference frame, *d* is the absolute difference, *w*(*p*) is the pixel’s contribution weight and *D* is the weighted mean pixel difference. RW computes the mean difference between a frame and every rest frame within a 3 s bidirectional window:

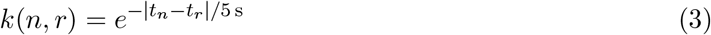

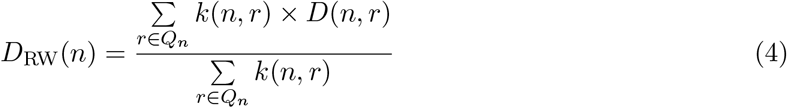

where *Q*_*n*_ is the set of reference frames within the window around frame *n*, excluding *n* itself, *t*_*n*_ and *t*_*r*_ are the times of frames *n* and *r*, and *k* is the relevance weight attributed to each reference, so that nearer references count more. RW works in three main steps (Fig. 1). First, a time-difference pre-pass with a 150 ms interval, chosen to exaggerate motion, finds the dataset’s highest-motion frames. Combining them, RW excludes pixels that do not contribute to motion and weights the rest (*w*(*p*) in Eq. 2) by their relative mean and constancy. A 20 ms time-difference pre-pass then locates a starting point, a quiescent portion of the dataset. Second, starting at that point, RW rolls through the recording, comparing each frame with the references accepted thus far and flagging new references when their amplitude and rates of change fall to rest levels. It then decides which frames should serve as references through a technique we name the Jury Consensus of Non-Motion Trend Observations (JCNTO, pronounced “Jacinto”; Fig. 1, step 6), in which independent detectors each decide where the data show rest, defining references when they reach consensus. Each detector on its own has weaknesses, but together they provide a more robust decision. Third, RW computes *D*_RW_ (Eq. 4) by comparing each frame with all of its local references. Because each frame is compared only with nearby references, run time remains linear; RW runs on a graphics processing unit (GPU), measures every region of interest in a field in one run, and uses subsampling and memory-mapped streaming where possible to substantially improve processing speed.

**Figure 1.**
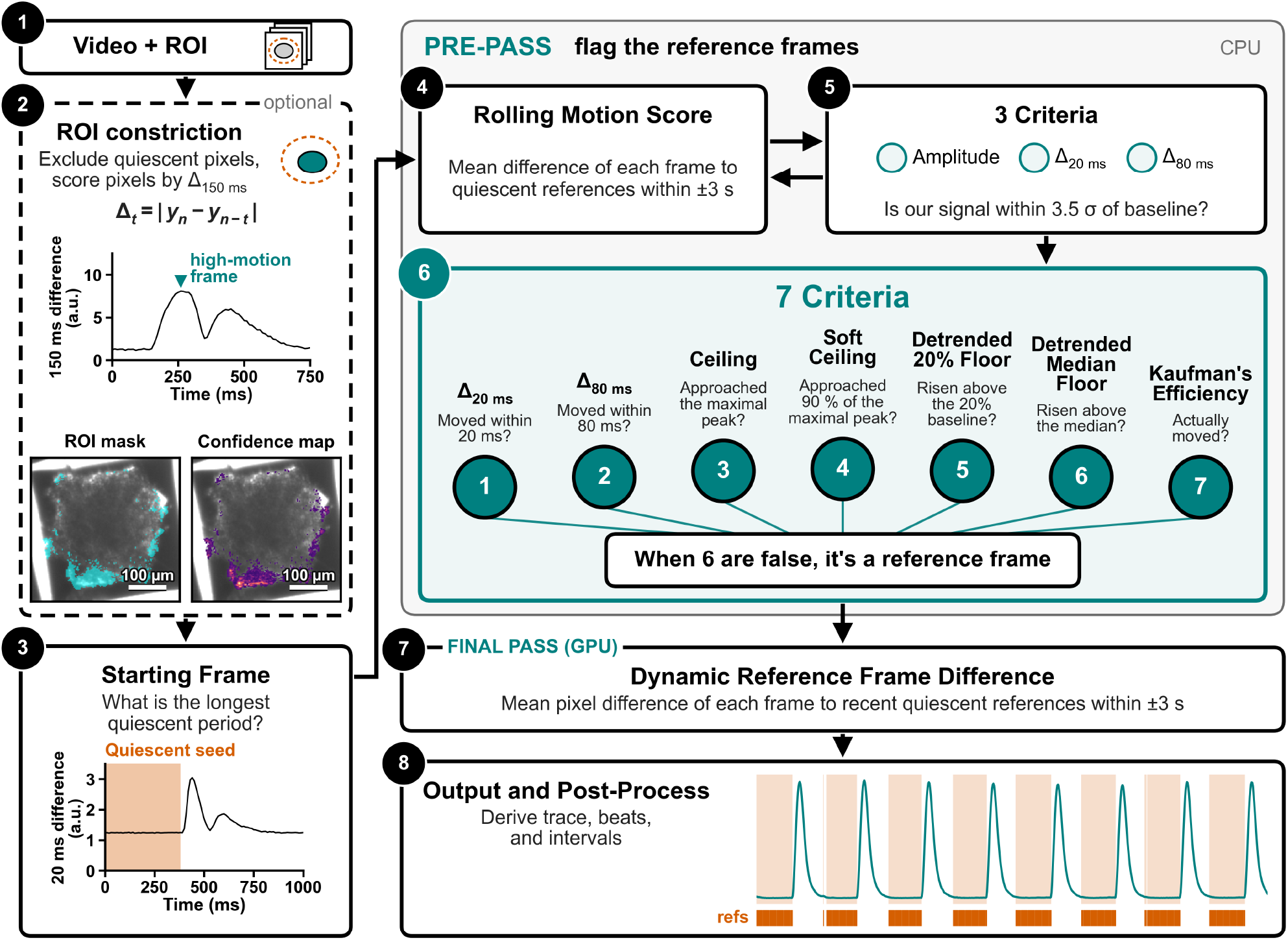
RhythmWriter compares motion relative to local recovery periods. **Step 1**, The user inputs an image sequence and defines one or more regions of interest (ROIs). **Step 2**, The ROI is constricted to exclude non-moving pixels, and each remaining pixel is weighted by its relative contribution, from the mean and constancy of its motion. **Step 3**, The pre-pass starts at the longest recovery period in the recording. **Steps 4 and 5**, The pre-pass flags whether each frame falls close to baseline. **Step 6**, The JCNTO voters define where the reference frames are in the sequence. **Step 7**, Feeding these reference frames to the graphics processing unit (GPU), RW computes the weighted mean difference between each frame and all local rest frames. **Step 8**, The data are output, and preliminary beats and intervals are defined.

The ROI constriction kept 1716 of the 17,956 pixels in its region of interest (9.6%; Fig. 1, step 2; Fig. 4b, c). The quiescent seed was located at 1327 s (step 3). JCNTO flagged 114,878 of the 193,868 frames (59%) as references (step 6), and the main output returned 1231 beats (step 8; Fig. 4d). Analyzing this recording took 33.5 s.

### RhythmWriter Validation

To validate RW’s output, we simulated a collection of 2000 rings with 5% opacity scattered around the centre of a 100 *×* 100-pixel field (Fig. 2a). With programmable displacement, interval and frequency, these emulate an aggregate, in which the outer rings displace more than the central ones. Gaussian noise with a standard deviation of 2% of the 8-bit range was added to every image to simulate camera grain. From the output complexes we derived each beat’s amplitude, inter-beat interval, recovery period and contraction–relaxation interval, defined as the time a beat spends above 10% of its amplitude (CRI90; Fig. 2b). With strength and CRI90 fixed, shortening the basic cycle length (BCL) from 1000 to 260 ms in 10 ms steps (Fig. 2c) left all 76 beats detectable, with every interval within 2.6% of the programme (Fig. 2f) and every amplitude within 2.1% of the first beat. Ramping the contractile strength from 100% down to 0%, up to 200% and back to 100% in 5% increments (Fig. 2d), RW transcribed all 80 contractions, and amplitude rose monotonically with strength with a median error of 6.0% (Fig. 2g). This non-linearity is already present in the simulated images, whose mean pixel change, without noise, read 25% strength as 56% and 200% as 152%; repeating these experiments on a hard-edged square, as a validation of edge-displacement measurement, gave linear amplitudes with a median error of 0.2% (Supplementary Fig. S1). Lengthening the programmed CRI90 of each beat’s smooth step twitch from 40 to 1528 ms at a fixed 500 ms recovery period and strength (Fig. 2e), RW assessed all 38 beats and derived CRI90 within 1.2% error at every step of 121 ms or longer (Fig. 2h), with amplitude within 4.6% of the first beat.

**Figure 2.**
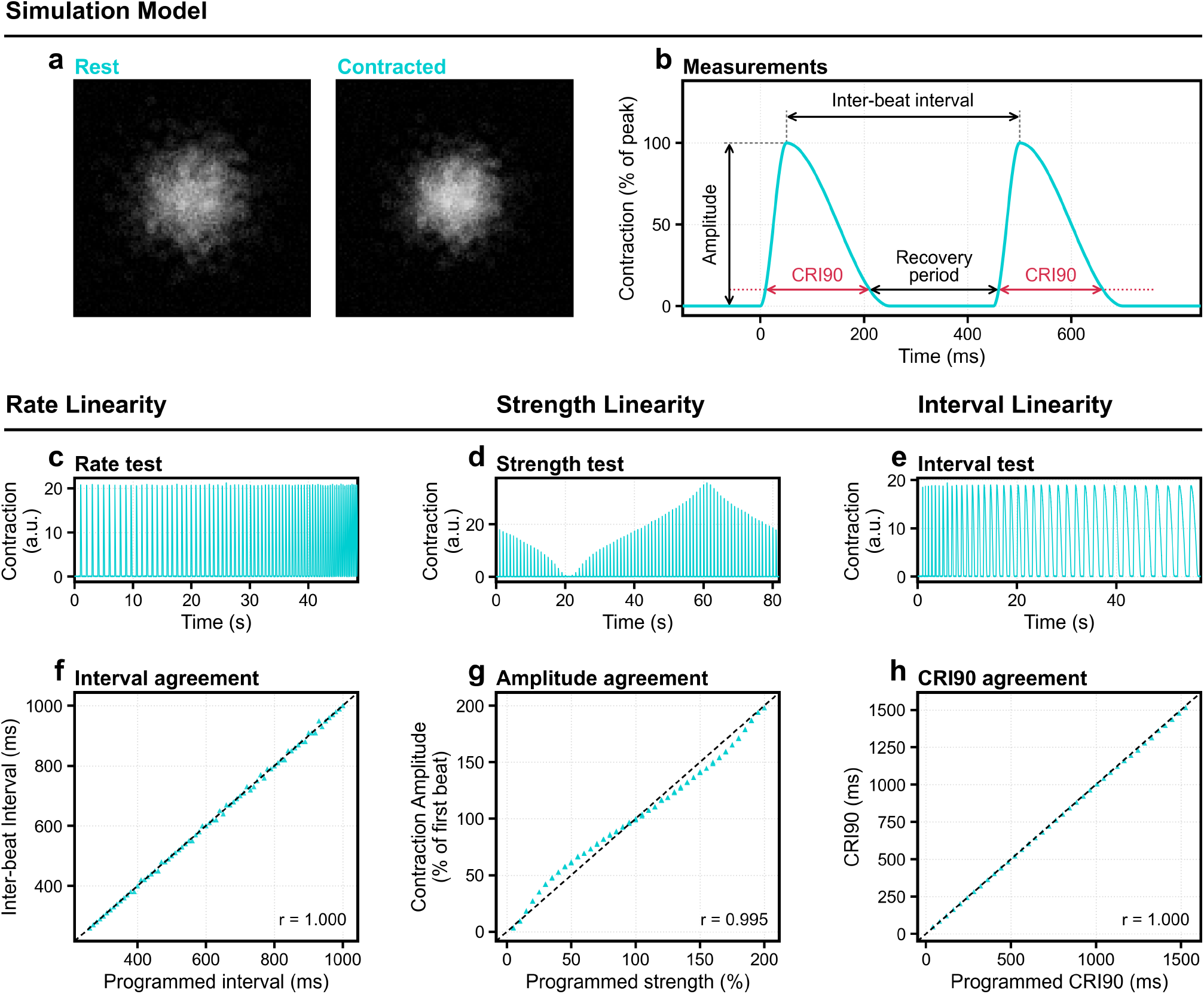
RhythmWriter detects intervals and amplitudes accurately and is robust to fast pacing. **a**, The simulated aggregate, 2000 rings of 5 % opacity in a 100 *×* 100-pixel field with Gaussian noise of standard deviation 2 % of the 8-bit range, at rest and at the peak of a 100 % beat. **b**, Two programmed twitches and the quantities measured. Amplitude is the height of the peak above rest. CRI90 (red) is the time a beat spends above 10 % of its amplitude. The inter-beat interval runs from peak to peak, and the recovery period is the inter-beat interval minus the CRI90 of the first beat. **c**–**e**, RhythmWriter’s output through three sweeps. **c**, Rate test, 76 beats with the programmed interval shortened from 1000 to 260 ms in 10 ms steps. **d**, Strength test, 81 beats whose programmed strength falls from 100 to 0 %, rises to 200 % and returns to 100 % in 5 % steps. **e**, Interval test, 38 beats whose programmed CRI90 lengthens from 40 to 1528 ms, with 500 ms of rest between beats. **f** –**h**, RhythmWriter’s measurements against the programmed values. **f**, Inter-beat interval (75 intervals). **g**, Amplitude as a percentage of the first beat (80 beats). **h**, CRI90 (38 beats). Dashed lines, identity. *r*, Pearson correlation. CRI90, contraction–relaxation interval at 90 % relaxation. a.u., arbitrary units.

RW’s outputs were validated for frame-rate invariance by assessing paired samples whose frame rate was altered by subsampling and frame interpolation, including non-standard sample rates. The mean Lin’s concordance with the native 100 frames per second (fps) was 0.997 (0.991–0.999; Supplementary Fig. S2). To validate start-time invariance, we compared the 8000-frame set to its 500-frame subdivisions. The relative amplitudes of contractions show marginal differences, caused by differences in each subdivision’s ROI, but remained comparable with a mean CCC of 0.98 (Supplementary Fig. S3). To validate noise robustness, randomized Gaussian noise was applied to our image set, and failure to capture beats only appeared once the noise reached 10% of the 8-bit range (Supplementary Fig. S4).

### RhythmWriter Computational Speed and Drift

Using an 8000-frame (80 s) stretch of the 32 min S1S2 recording of Fig. 4, we ran paired tests against MM (Fig. 3a). In this process we found that a bug in their noise-reduction step, which excludes non-moving pixels, made its run time grow cubically with the number of frames, so that it took 3.1 h on average to compute the 8000 frames. Without that noise reduction, the drift degradation worsened so all experiments were performed with it on. RW was initially 62 times faster on the shortest recording, with a start-up time of 6 ms against 1.06 s and became comparatively more efficient as a function of frames. MM’s baseline drifted away from its reference frame by 0.61% of full scale per second, while RW’s did not (Fig. 3b). We placed the reference frame of MM, near the start and near the end of the recording and compared the amplitudes and CRI90s of their paired beat complexes (Fig. 3c, d). The outputs disagreed, with amplitudes differing by a median of 13% (up to 36%) and CRI90s by a median of 6% (up to 21%), where the amplitude disagreement grew with the difference between the two outputs’ baselines (*r* = 0.99), from a median of 1% midway between the reference frames to 18% and 35% at the start and end of the recording.

**Figure 3.**
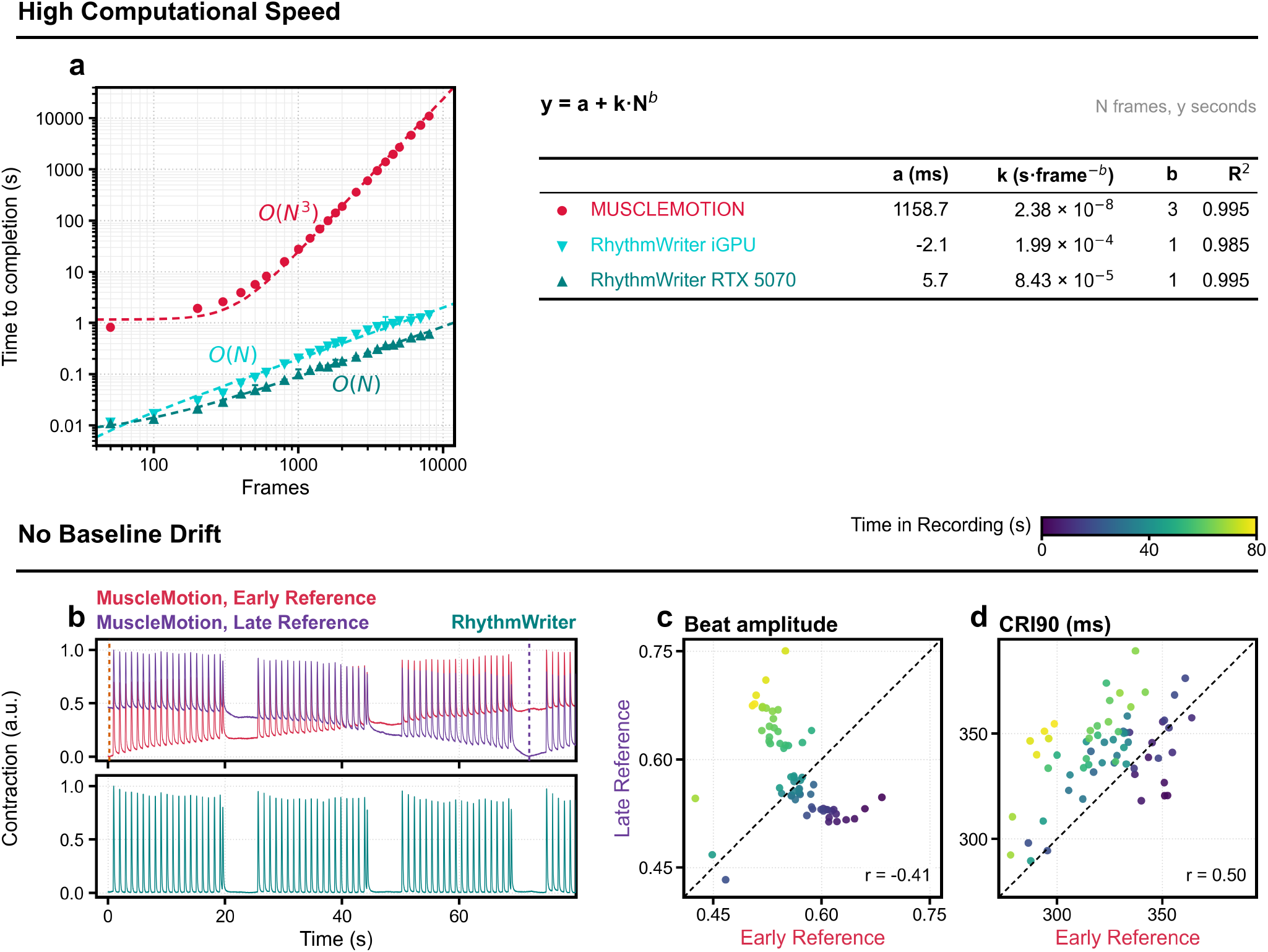
RhythmWriter is fast and drift-immune. **a**, Run time against the number of frames analyzed (50–8000 frames of the 80 s test recording) for MUSCLEMOTION and for RhythmWriter on an integrated GPU (iGPU) and an RTX 5070. Points, median of three runs. Bars, range. Dashed lines, fits of *y* = *a* + *kN* ^*b*^. **b**, The 8000-frame recording analyzed by MUSCLEMOTION with an early (0.18 s, red) and a late (71.94 s, purple) reference frame, and by RhythmWriter (teal), each scaled from 0 to 1. Dashed lines, the reference frames. **c, d**, Amplitude (**c**) and CRI90 (**d**) of each beat in the two MUSCLEMOTION runs of **b** (64 beats), coloured by time in the recording. Dashed lines, identity. *r*, Pearson correlation.

**Figure 4.**
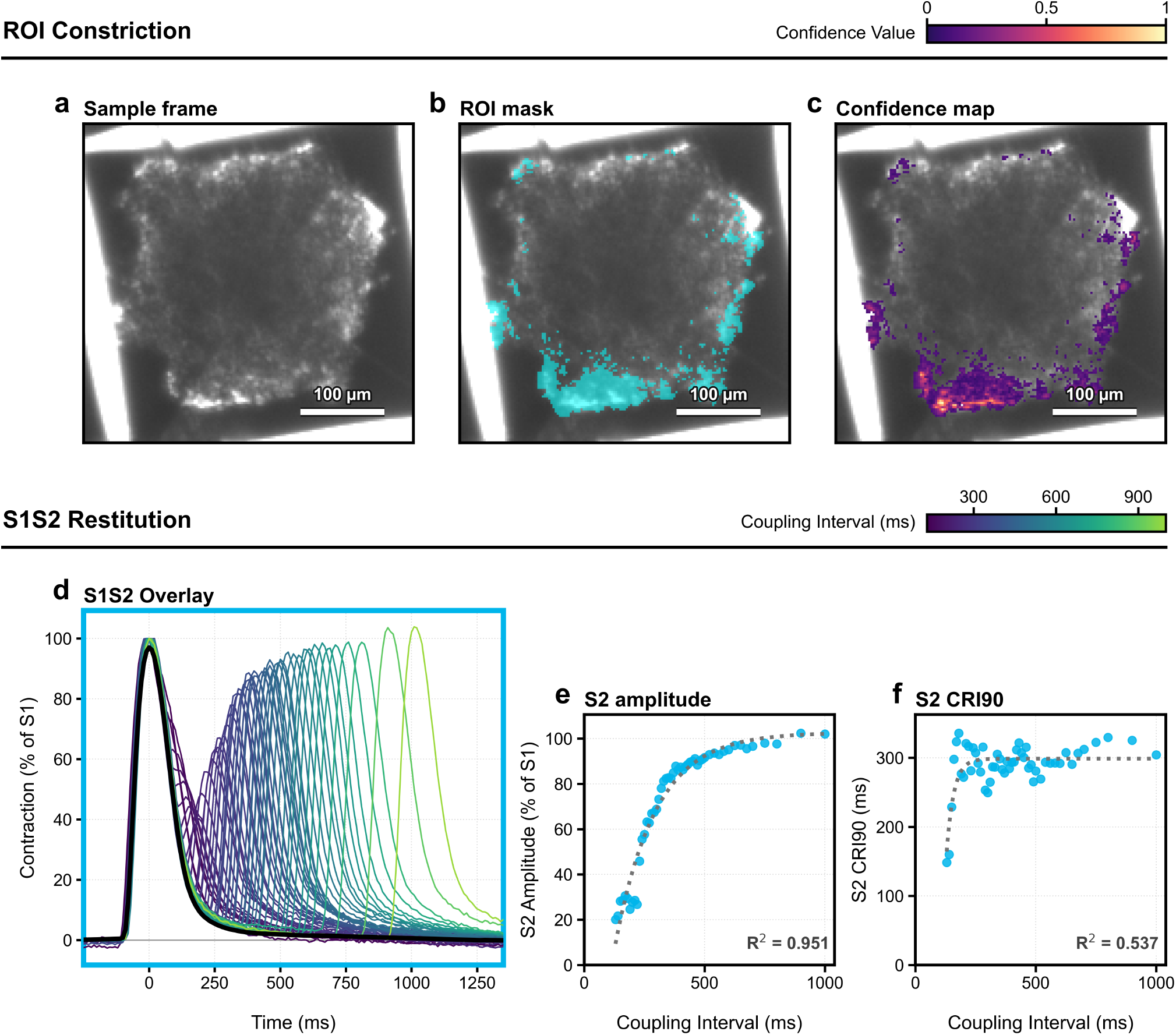
S1S2 non-invasive mechanical restitution. Aggregate 2 (Fig. 6a) under the S1S2 protocol. **a**, The first frame, cropped around the input ROI. **b**, The pixel set kept by the ROI constriction (cyan, 1716 of 17,956). **c**, Per-pixel contribution weights (confidence), scaled from 0 to 1. Scale bars, 100 *µ*m. **d**, The 19th S1 and the S2 at each of the 51 couplings (130–1000 ms), aligned on the S1 peak and scaled to it. Colour, coupling interval. Black, median S1. **e**, S2 amplitude as a percentage of the 19th S1. **f**, S2 CRI90. Dotted lines, exponential fits with their *R*^2^ (time constant 168 ms in **e**). S1, conditioning beat. S2, premature beat. Blue, aggregate 2 of Fig. 6.

### Mechanical Restitution

#### S1-S2 Protocol

In the same recording, aggregates were paced for 25.7 min with an S1S2 protocol of 19 S1 stimuli at a BCL of 1000 ms, followed by a stimulus of increasing prematurity (S2), from 1000 ms down to 10 ms, and a period of 6 s of rest (Fig. 4d). Each S1 train began with a recovery from rest, in which the first S1 after its 6 s of rest was 9.7% stronger and its CRI90 25 ms longer than in the steady state (Supplementary Fig. S5). Averaged over the 47 trains, amplitude settled within 1% of its steady state by the fourth beat and CRI90 by the third, at a steady state of 8.8 a.u. and 260 ms. The 51 S2s at coupling intervals of 130 ms or longer were measured by RW. S2 amplitude was 82–102% of the S1 at couplings from 350 through 1000 ms and then fell steeply to 20% at 130 ms (Fig. 4e), recovering with a single time constant of 168 ms (*R*^2^ = 0.951). S2 CRI90 stayed between 249 and 336 ms at couplings of 160 ms and above, shortening sharply below that (Fig. 4f).

#### Dynamic Protocol

In this same aggregate, a dynamic ramp protocol was run for 12.1 min, ramping from 60 to 245 bpm (BCL 1000 to 245 ms) in 62 steps of 20 stimuli (Fig. 5a). The aggregate followed 1:1 entrainment throughout, answering all 1240 stimuli with a beat (Fig. 5b, c). As the recovery time, the cycle length minus the preceding beat’s CRI90, shortened from about 750 to 80 ms, contraction amplitude fell by 82% (Fig. 5d) and CRI90 by 40% (Fig. 5e), the latter with a time constant of 201 ms.

**Figure 5.**
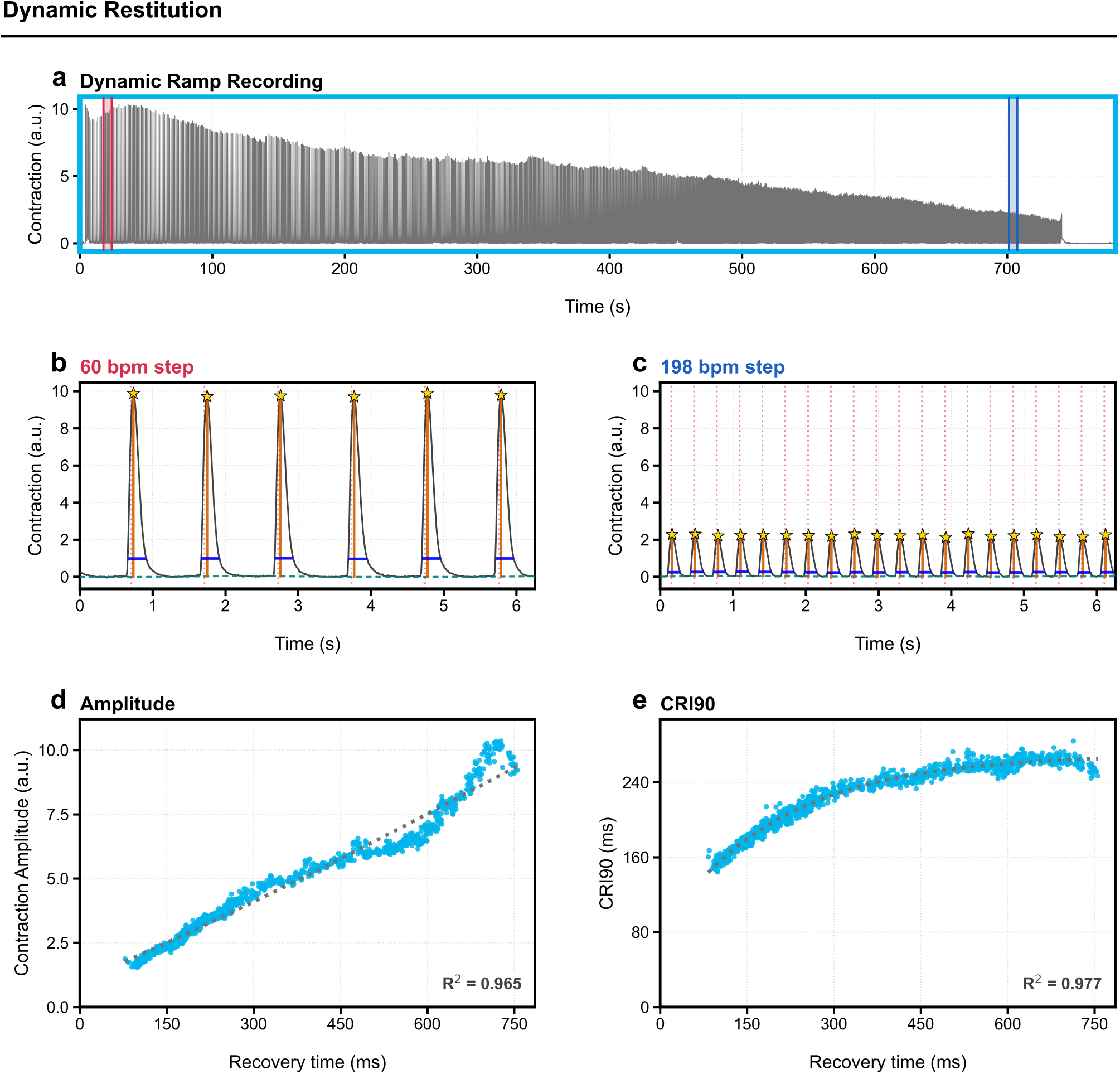
Dynamic ramp mechanical restitution. Aggregate 2 (Figs 4 and 6) under dynamic pacing, 62 steps of 20 stimuli as the programmed cycle length shortened from 1000 to 245 ms. **a**, The whole recording (13.0 min), the last 40 s unpaced. Red and blue, the windows of **b** and **c. b, c**, The last 6 s of the 60 bpm step (1000 ms; **b**) and the 198 bpm step (303 ms; **c**), on one scale. Grey, the trace. Dashed teal, its baseline. Dotted red lines, the stimulus schedule. Stars, beats. Orange, amplitude. Blue, CRI90. **d, e**, Amplitude (**d**) and CRI90 (**e**) of each beat against its recovery time, the cycle length minus the preceding beat’s CRI90. Dotted lines, exponential fits with their *R*^2^ (time constant 201 ms in **e**). Blue, aggregate 2 of Fig. 6.

#### High Throughput S1-S2 restitution

To demonstrate scalability, RW assessed the motion across five aggregates paced in parallel within a single field of view (Fig. 6a). Each yielded their own restitution plots, similar to the ones shown in Figs. 4 and 5 (Fig. 6b). Each aggregate produced S2 beats at all 51 coupling intervals from 130 to 1000 ms. Their restitution curves differed (Fig. 6c), showing that functional assessment can be derived on a per-aggregate basis. In total, this took 170.8 s for 161.6 min worth of data across the five aggregates (34.2 s per aggregate).

**Figure 6.**
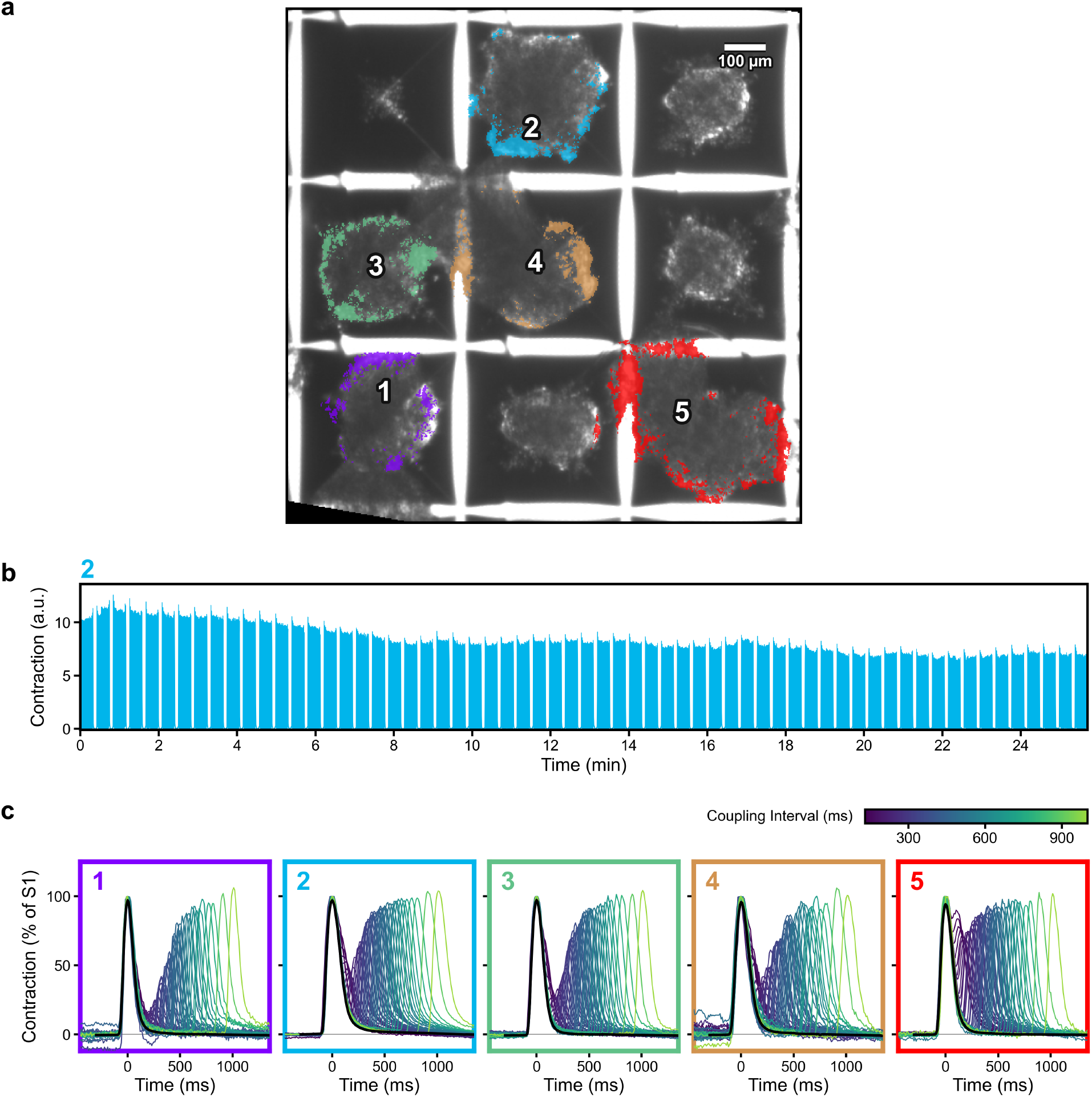
RhythmWriter enables analysis of long datasets across several samples. Five aggregates in one camera field under the S1S2 protocol, measured in one RhythmWriter run. **a**, The field, rotated to align the microwell walls and cropped to the nine microwells around the five aggregates, with the pixels kept by each aggregate’s ROI constriction in colour. Black, outside the camera field. Scale bar, 100 *µ*m. **b**, Aggregate 2 over the whole S1S2 protocol (25.7 min). **c**, The 19th S1 and the S2 at each of the 51 couplings (130–1000 ms) for each aggregate, as in Fig. 4d. Colour, coupling interval. Black, median S1. Frame colour, the aggregate in **a**.

## Discussion

RW improves upon previously established methods by achieving time-scale invariance and drift immunity, circumventing the bias of a single reference frame. This lack of drift makes contractility comparable across long datasets (Fig. 3b), and RW proves robust at fast pacing frequencies (Fig. 2c, f). On the recordings published with MUSCLEMOTION, RW abolished baseline drift across five tissue types (Supplementary Fig. S6). PLoRI, another published approach to drift immunity, compares each frame with a rolling median [28]. Its output degraded as recovery intervals shortened, making it unsuitable for fast pacing (Supplementary Fig. S7). Together with its gains in computational speed, RW provides a framework for the scaled analysis of hiPSC-CM contractility. To our knowledge, this is the first mechanical restitution experiment derived non-invasively from hiPSC-CMs under a paired, high-resolution S1S2 and dynamic protocol (63 coupling intervals and 62 pacing steps), where all beats were resolved. Label-free video also has advantages over calcium or voltage dye imaging, since several aggregates can be monitored through 45 min of recording without the phototoxicity and photobleaching that make such dyes unsuitable for long-term measurements [37, 44]. This allows numerous pacing protocols to be performed in succession, providing finer detail for functional models on a per-aggregate basis.

Restitution across the five aggregates showed that RW resolves differences in contractility dynamics on a per-aggregate basis (Fig. 6c). Under dynamic pacing, 1:1 capture held as the BCL shortened to 245 ms, meaning the refractory period of the aggregates remained below that cycle length. Further shortening of the BCL would be expected to cause a loss of 1:1 capture. With S2 beats persisting down to 130 ms in all five aggregates, their refractory period after a 60 bpm steady state was at most 130 ms. RW’s amplitude represents a relative measure of contractile motion, a proxy for contractile force [21, 22]. As mechanical restitution has been described to reflect calcium cycling [29, 30], which is itself dependent on the cardiac action potential [23, 45], alterations to any of the components that contribute to contraction can be expected to appear in these restitution dynamics. Should that prove true, this would make mechanical restitution a promising candidate for an easy hiPSC-CM follow-up study that can complement drug profiles [11].

RW’s output amplitudes are relative, meaning comparisons can only be made within a single run. Normalized protocols, such as our restitution protocols, make these dynamics comparable because they measure relative changes. This is fundamentally the biggest limitation of the algorithm, since the relative output signal depends on the region of interest (Supplementary Fig. S3). To capture the tissue’s full range of motion and limit the signal saturation others have described [21], our ROI is defined from a sample of high-motion frames, one per 20 s, which normalizes the scale of motion relative to the strongest beat of the recording. RW also cannot accurately resolve CRI90s for contractions that are brief relative to its sampling rate. This is why our data were captured at 100 fps. RW also presumes that there is a good reference frame within *±*3 s of any given frame.

Regulators have expressed a growing interest in human in vitro cardiomyocyte data [11, 12]. With RW, one well can yield high-resolution data from several aggregates in parallel within a single dataset. This is an experiment that any laboratory with a microscope, a camera, a field stimulator and a computer can perform, and an integrated GPU is sufficient to run the algorithm (Fig. 3a). Because no specialized preparation is needed, patient-derived lines can be compared through protocols such as these. The length of a protocol and the number of aggregates now become choices for the researcher, rather than limitations of the technique.

## Methods

### Cell line and ethics

Human induced pluripotent stem cells (hiPSCs) of the line STAN248i-617C1 (WiCell; RRID: CVCL_JS07; female donor aged 38 years), reprogrammed from peripheral blood within the Next Generation Genetic Association Studies program, were used [46]. In this wild-type line, CRISPR– Cas9 genome editing had previously inserted an extracellular hemagglutinin (HA) tag homozygously between residues 443 and 444 of hERG (exon 6 of *KCNH2* ) for a separate study of hERG trafficking [47]. This tag was not used in this study. The Stanford Institutional Review Board approved the donor study protocol, and the donor gave written informed consent [46]. All methods were carried out in accordance with relevant guidelines and regulations.

### hiPSC culture

hiPSCs were maintained on hESC-qualified Matrigel (Corning, 354277) in mTeSR Plus medium (STEMCELL Technologies, 100-0276) at 37 °C and 5% CO_2_, with daily medium changes. The cells were washed with DPBS (Gibco, 14190250) and passaged every 3–4 days at a 1:20 ratio with ReLeSR (STEMCELL Technologies, 100-0483) to minimize spontaneous differentiation. Passaged cells were replated in mTeSR Plus supplemented with 10 *µ*M Y-27632 (STEMCELL Technologies, 72304). Each culture was passaged no more than 10 times before a new vial was thawed. Cultures were tested weekly for mycoplasma, which was never detected. After genome editing, more than 98% of the HA-tagged hiPSCs expressed the pluripotency markers SSEA-4 and TRA-1-60 by flow cytometry [47].

### Cardiomyocyte differentiation and purification

hiPSCs were differentiated into ventricular hiPSC-CMs by temporal modulation of Wnt signaling, adapted from Lian et al. [48] hiPSCs were seeded at 200,000 cells per well of a 6-well plate so that they reached confluency quickly. When the cells reached 80% confluency (day 0), the medium was replaced with RPMI 1640 (Gibco, 11875119) supplemented with 2% B-27 minus insulin (Gibco, A1895601) and 12 *µ*M CHIR99021 (STEMCELL Technologies, 72054) for 24 h. IWP-4 (5 *µ*M, STEMCELL Technologies, 72554) was added on day 3 in a 1:1 mix of fresh and used medium and removed on day 5. From day 7, the cells were fed every 48 h with maintenance medium consisting of RPMI 1640 with 2% B-27 Plus (Gibco, A3582801). Spontaneous beating began by day 9 and spread throughout the monolayer by day 12. hiPSC-CMs were then enriched by metabolic selection [49] for 3 days in glucose-free RPMI 1640 (Gibco, 11879020) with 2% B-27 (Gibco) and 5 mM sodium L-lactate (Sigma-Aldrich, 71718) and returned to maintenance medium.

### Aggregates and wells

hiPSC-CMs were washed with PBS (Gibco, 10010049) and dissociated in two steps, each with 1 mL per well for 10 min at 37 °C and 5% CO_2_. The first step used 0.1% collagenase type II (Gibco, 17101015) in RPMI 1640, and the second used TrypLE Express (Gibco, 12605028). Each well then received 1 mL of maintenance medium with 10 *µ*M Y-27632, and the cells were collected, centrifuged for 4 min at 290 *× g* and counted. AggreWell400 plates (STEMCELL Technologies, 34415) were pretreated with Anti-Adherence Rinsing Solution (STEMCELL Technologies, 07010), seeded with about 1.2 million live cells per well (about 1,000 per microwell) and centrifuged at 100 *× g*, following the manufacturer’s instructions. The aggregates were cultured in maintenance medium and recorded in their microwells 7 days after aggregation.

All recordings come from one well of a single differentiation, treated with 0.1% dimethyl sulfoxide (DMSO). The same camera field was recorded under both pacing protocols. One region of interest (ROI) was drawn around each aggregate in the field, and non-contracting tissue was excluded. All five ROIs were measured in a single RW run.

### Electrical pacing

Prior to recording, the medium was replaced with RPMI 1640 with HEPES (Gibco, 22400089) supplemented with B-27 Plus, and aggregates were held at 37 °C. Aggregates were field-stimulated through a C-Dish 24-well carbon-electrode insert driven by a C-Pace EM stimulator (IonOptix), triggered by an Arduino. Pulses were biphasic, 10 ms long and 40 V. The S1S2 protocol consisted of 63 trains of 19 S1 stimuli at 1000 ms, each followed by an S2 and 6 s of rest. The S2 coupling interval shortened from 1000 to 10 ms in steps of 100 ms (to 800 ms), 50 ms (to 700 ms), 25 ms (to 600 ms), 20 ms (to 500 ms) and 10 ms (to 10 ms). The dynamic protocol shortened the pacing interval from 1000 to 245 ms over 62 steps of 20 stimuli, without pause.

### Video recording

Aggregates were imaged in their microwells by phase-contrast microscopy on a Zeiss Axiovert 135 inverted microscope with a Plan-Neofluar 2.5*×*/0.075 objective with a Basler acA720-520um camera, at 100 fps using pylon Viewer, as 8-bit TIFF image sequences of 728 *×* 540 pixels. The dynamic recording (13.0 min) was performed first, followed 25 s later by the S1S2 recording (32.3 min).

### Beat measurements

Beats were detected on RW’s output using a Python script, as local maxima at least 0.18 s apart whose prominence exceeded six times the local noise and 35% of their height above the diastolic floor. Each beat was measured against a local baseline drawn through the minimum of each inter-beat interval. Amplitude is the peak above this baseline, and CRI90 is the time between the crossings of 10% of the amplitude on either side of the peak, interpolated between frames. CRI90 was left unmeasured when either crossing fell beyond a neighboring beat. Under dynamic pacing, each stimulus was assigned the highest peak within half a cycle of its scheduled time that rose at least 0.5 a.u. above the preceding trough.

### Simulations

Simulated stacks (Fig. 2, Supplementary Fig. S1) were 8-bit, 100 *×* 100 pixels at 100 fps. The aggregate was 2000 rings of 5% opacity and the square a hard-edged 40 px square, each contracting about the field centre with a swept area proportional to programmed strength. Twitches rose over 20% and relaxed over 80% of their duration, and Gaussian noise of 2% of the 8-bit range was added to every frame. The rate, strength and duration sweeps held 76, 81 and 38 beats. A beat was found when a detected peak lay within 150 ms of its programmed peak. The noise test (Supplementary Fig. S4) used the strength sweep of an earlier simulation of 2000 solid discs, with noise of 1 to 150 grey levels.

### Validation corpus

RW’s bidirectional window, *τ* and weighting constants were chosen through iterative assessment and validation on a corpus of 85 test recordings (simulations, negative controls, aggregates and real tissue), designed to test specific features that contribute to the signal-to-noise ratio.

### MUSCLEMOTION Analysis

MUSCLEMOTION v1.0 was run unmodified in Fiji using default settings. Reference frames were selected automatically, or set manually (Fig. 3b, 71.94 s). Baseline drift is the change in a trace’s median rest level from before the first beat to the last pause, per second, as a percentage of the trace’s range. The two reference runs were compared beat by beat by the relative difference in amplitude and CRI90. Run time (Fig. 3a) was measured on the first 50 to 8000 frames of the test recording, three times each, and fitted as *y* = *a* + *kN*^*b*^, with *b* fixed at 3 for MUSCLEMOTION and 1 for RhythmWriter (free fits gave 2.80 and 0.95–1.10). RW was timed on an NVIDIA RTX 5070 and an integrated Intel UHD 770 GPU, on one workstation (Intel Core i7-14700K, 32 GB RAM).

### RhythmWriter Analysis

Each recording was analyzed in a single RW run, measuring each ROI in sequence. Run time per ROI was logged by the plugin and excludes starting Fiji and opening the image sequence.

### Restitution analysis

Each S1S2 train comprised 19 S1 beats and an S2. Trains were screened by eye, and the 47 trains of aggregate 2 kept at couplings of 170–1000 ms were averaged for recovery from rest. S1 amplitude was measured above the train’s diastolic level, each train’s steady state was the median of its beats 5–18, and the first S1 was compared with that steady state as the median over trains. To isolate the S2, a template of the aggregate’s S1 beat was fitted to the 19th S1 and subtracted with the S1’s exponential decay, leaving a residual. An S2 was detected when the residual exceeded 3.5 robust standard deviations of its pre-S2 level. S2 amplitude, as a percentage of the 19th S1, and S2 CRI90 were measured on this residual at the 51 couplings from 130 to 1000 ms. Under dynamic pacing, recovery time was the programmed cycle length minus the CRI90 of the preceding beat. Recovery curves were fitted by least squares with one or two exponentials, and time constants are reported only for fits whose parameters did not lie at their bounds (Figs 4e and 5e).

### Statistics

Error bars (Supplementary Fig. S5) are the standard error of the mean over the 47 trains of aggregate 2. The five aggregates came from one well of a single differentiation and are technical replicates, so no statistic was pooled across them. Correlations are Pearson’s *r*, and agreement between traces is Lin’s concordance correlation coefficient [50]. No hypothesis tests were performed.

### Software

RhythmWriter is written in Java and was run with Fiji 2.16.1 (ImageJ 1.54p) on Java 21.0.7. Analyses used Python 3.13.14 with NumPy 2.4.3, SciPy 1.17.1, Matplotlib 3.10.8 and pandas 3.0.2.

### Use of AI tools

Anthropic Claude was used in the implementation and coding phase of this program. Figures were generated with the help of Claude but have been validated by the authors.

## Data Availability

The datasets generated and analyzed during the current study are available from the corresponding author on reasonable request.

## Code availability

RhythmWriter is open-source and available at https://github.com/BrandonPerestrelo/RhythmWriter_GPU. The analysis scripts are available from the corresponding author on reasonable request.

## Acknowledgements

We thank Leon Glass for his help in preparing the manuscript, Marium Butt (Simon Fraser University, British Columbia) for generating the CRISPR-tagged hiPSC lines, and Fedgi Gaspard for support in culturing and differentiating hiPSC-CMs. This work was supported by the Canadian Institutes of Health Research (CIHR) and the Natural Sciences and Engineering Research Council of Canada (NSERC).

Naming of the JCNTO was in reference to the first author’s late grandfather, who having spent his entire life as the graveskeeper for the Mount Royal cemetery, would likely have learned to avoid hills where possible.

## Author contributions

B.P. conceived the project, wrote the RhythmWriter software and led the project. M.L. supported the culturing and differentiation of hiPSC-CMs and wrote the Arduino pacing-protocol script. A.S. and G.B. jointly supervised the work. All authors reviewed the manuscript.

