## Supplemental Figures for "RhythmWriter: GPU-accelerated Contractility Quantification for Large Datasets"

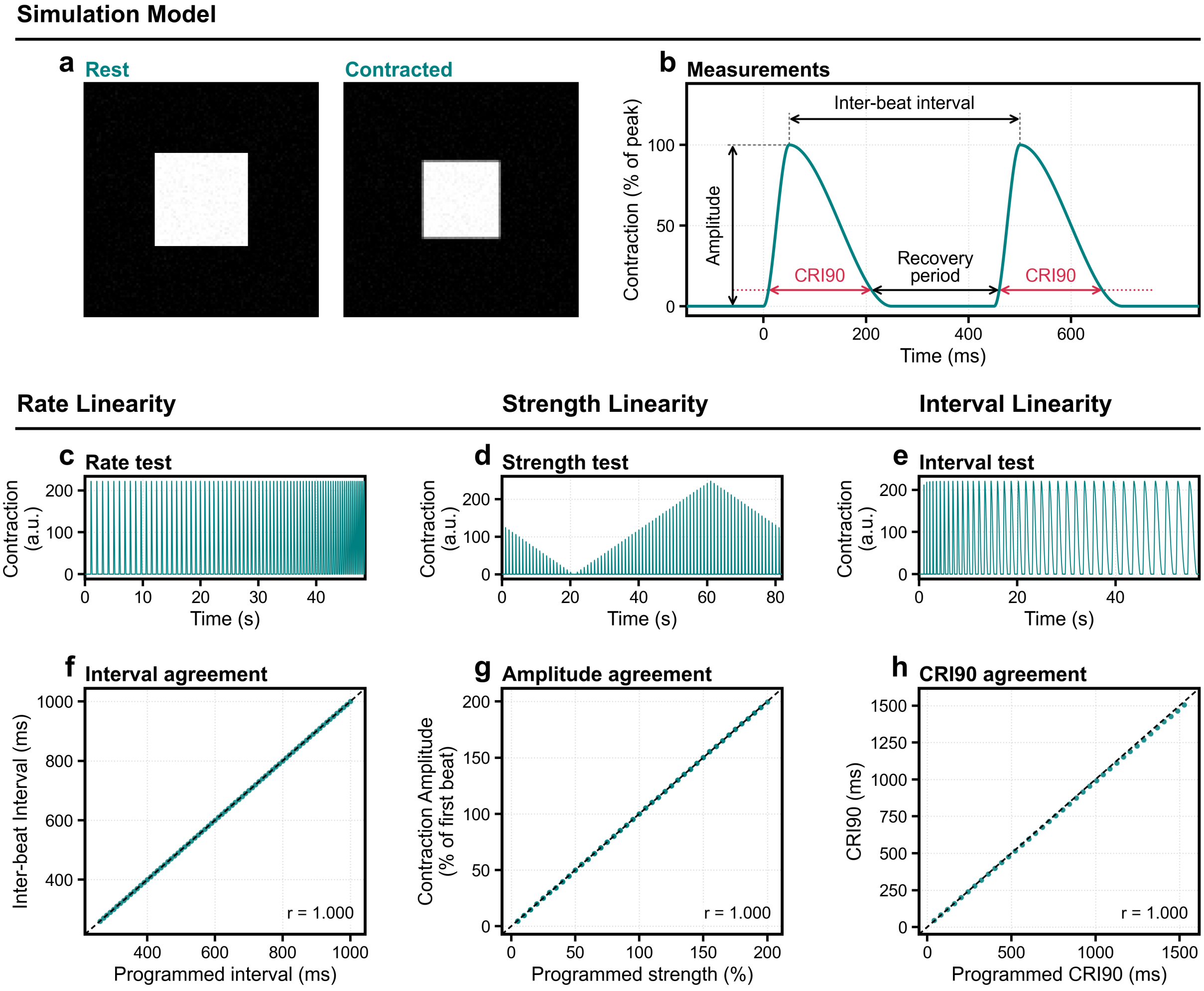


**Supplementary Figure S1| RhythmWriter detects intervals and amplitudes accurately in edge measurements.** **a**, The simulated square, hard-edged and 40 pixels wide in a 100 × 100-pixel field with Gaussian noise of standard deviation 2% of the 8-bit range, at rest and at the peak of a 100% beat. Its swept area is exactly proportional to programmed strength. **b**, Two programmed twitches and the quantities measured, as in Fig. 2b. **c**–**e**, RhythmWriter's output through the rate, strength and interval tests of Fig. 2. **f**–**h**, RhythmWriter's measurements against the programmed values. **f**, Inter-beat interval (75 intervals, all exact). **g**, Amplitude as a percentage of the first beat (80 beats, median absolute error 0.2 percentage points). **h**, CRI90 (38 beats, median error −1.2%). Dashed lines, identity. $r$, Pearson correlation.

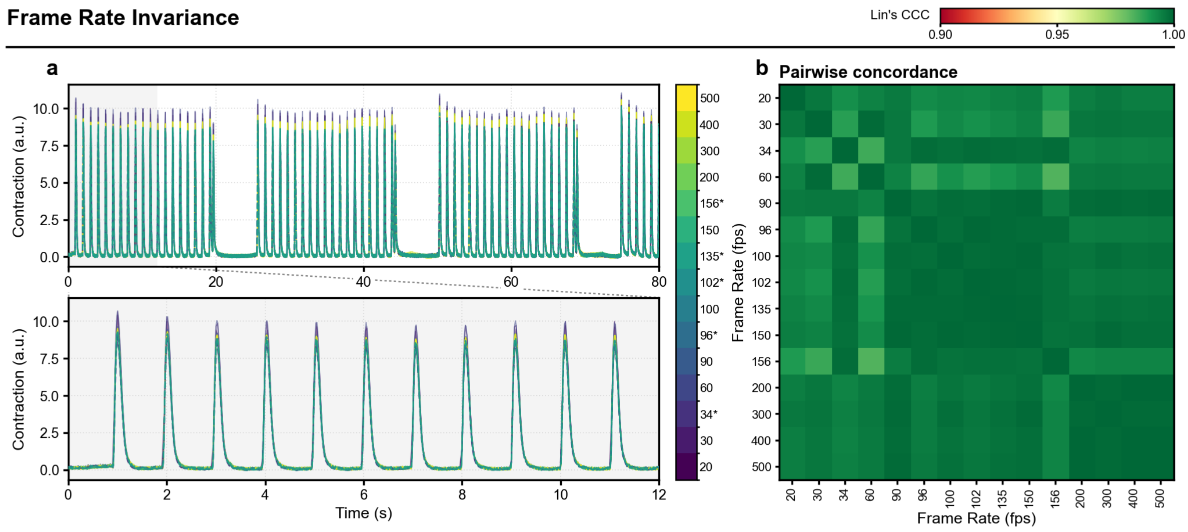


**Supplementary Figure S2| RhythmWriter is frame-rate invariant.** The 80 s test recording of Fig. 3 (8000 frames at 100 frames per second, fps) was resampled to 14 other rates from 20 to 500 fps, and RhythmWriter analyzed each version as a separate recording. **a**, The output at all 15 rates (colour), over the whole recording (top) and its first 12 s (bottom, the shaded span above). Asterisks, non-standard sample rates. **b**, Lin's concordance correlation coefficient (CCC) between the outputs at every pair of rates, after linear interpolation onto a common 100 Hz grid. Against the native 100 fps, the CCC averaged 0.997 (range 0.991–0.999). a.u., arbitrary units.

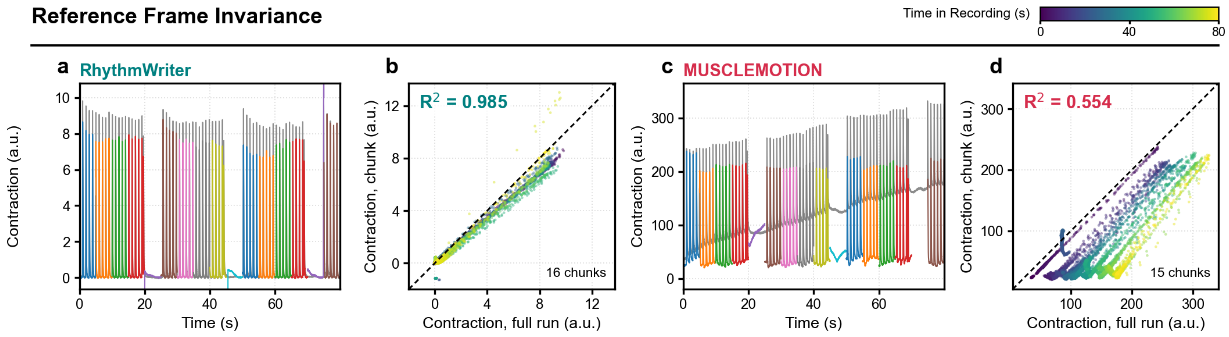


**Supplementary Figure S3| RhythmWriter is ROI-dependent.** The 80 s test recording of Fig. 3 was analyzed whole and as sixteen consecutive 500-frame (5 s) chunks, each chunk as a separate recording placed back at its time. **a**, RhythmWriter, the full run (grey) and the chunk runs (colours), each chunk with its own reference frames, ROI constriction and pixel weights. **b**, Every frame of the RhythmWriter chunk runs against the full run, coloured by time in the recording. Lin's concordance correlation coefficient over these points is 0.981. **c**, **d**, The same for MUSCLEMOTION, with 15 chunks (no MUSCLEMOTION output exists for the 70–75 s chunk). Dashed lines, identity. $R^2$, squared Pearson correlation. a.u., arbitrary units.

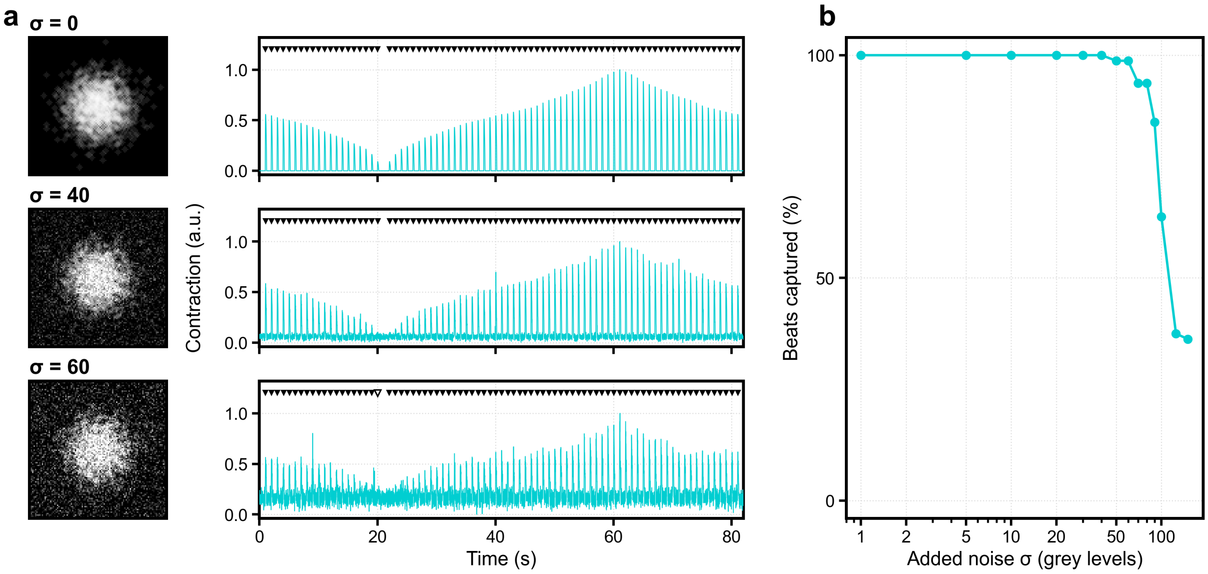


**Supplementary Figure S4| RhythmWriter is noise-resistant.** Gaussian noise added to the strength test of an earlier simulated aggregate, 2000 solid discs of 10% opacity in a 100 × 100-pixel field, with strength stepped from 100 to 0%, up to 200% and back to 100% (80 programmed beats). **a**, Rows at noise standard deviations $\sigma$ of 0, 40 and 60 grey levels. Left, a rest frame. Right, RhythmWriter's output, scaled from 0 to 1, with the programmed beats above (filled where found, open where lost). **b**, Percentage of the 80 programmed beats found against $\sigma$ (logarithmic axis). Every beat was found up to $\sigma = 40$ (15.7% of the 8-bit range), and the first beat lost, at $\sigma = 50$, was the weakest (5% strength). Capture at $\sigma = 125$ and 150 is at chance level.

### Recovery from Rest - S1 Train

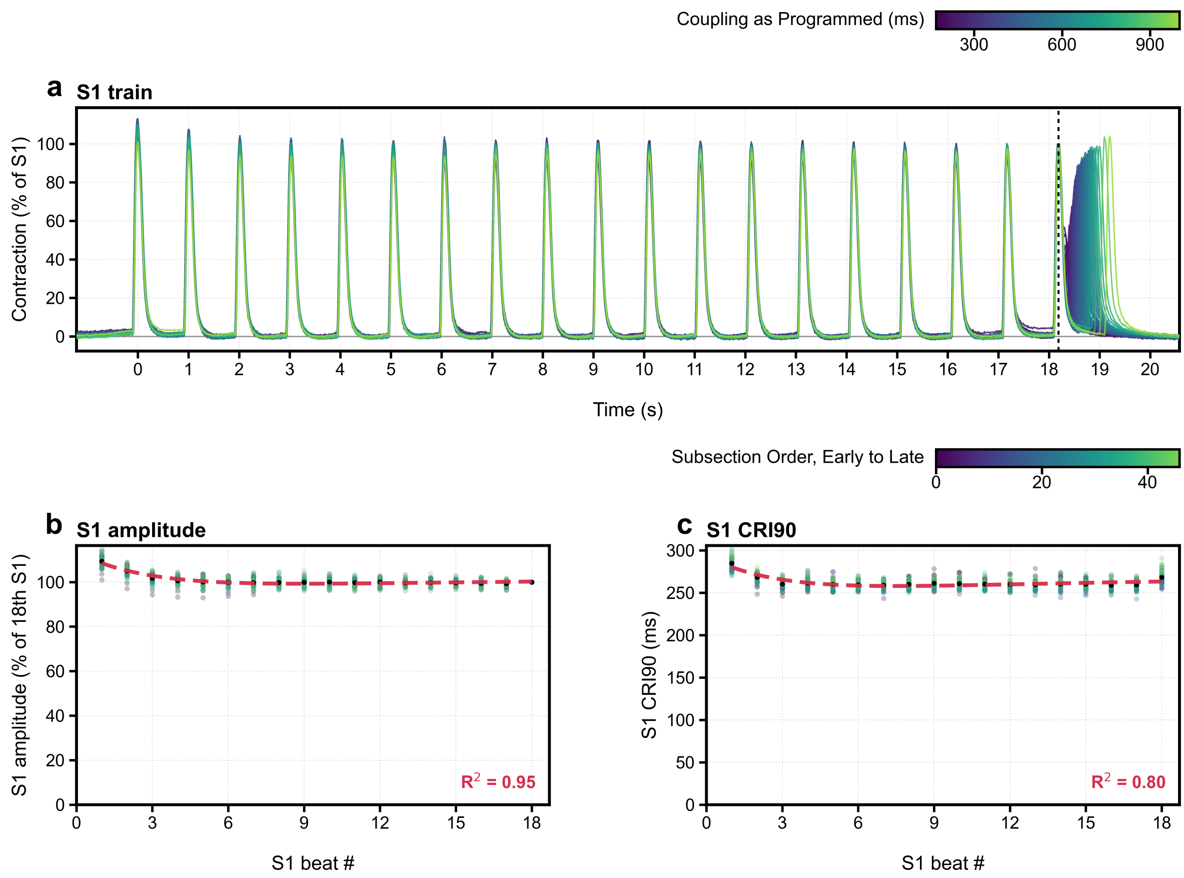


**Supplementary Figure S5| Recovery from 6 s of rest during the S1 train.** Aggregate 2 under the S1S2 protocol, over the 47 trains kept at couplings of 170–1000 ms (Methods). **a**, RhythmWriter's trace of each train, aligned on its 19th S1 (dotted line) and scaled to it. Colour, coupling interval. Time zero is the first S1. **b**, **c**, Amplitude (**b**) and CRI90 (**c**) of S1 beats 1–18, with amplitude as a percentage of the 18th S1. Faint points, each train, coloured by its order in the recording. Black, mean ± s.e.m. Dashed red, two-exponential fits to the means with their $R^2$. S1, conditioning beat. S2, premature beat.

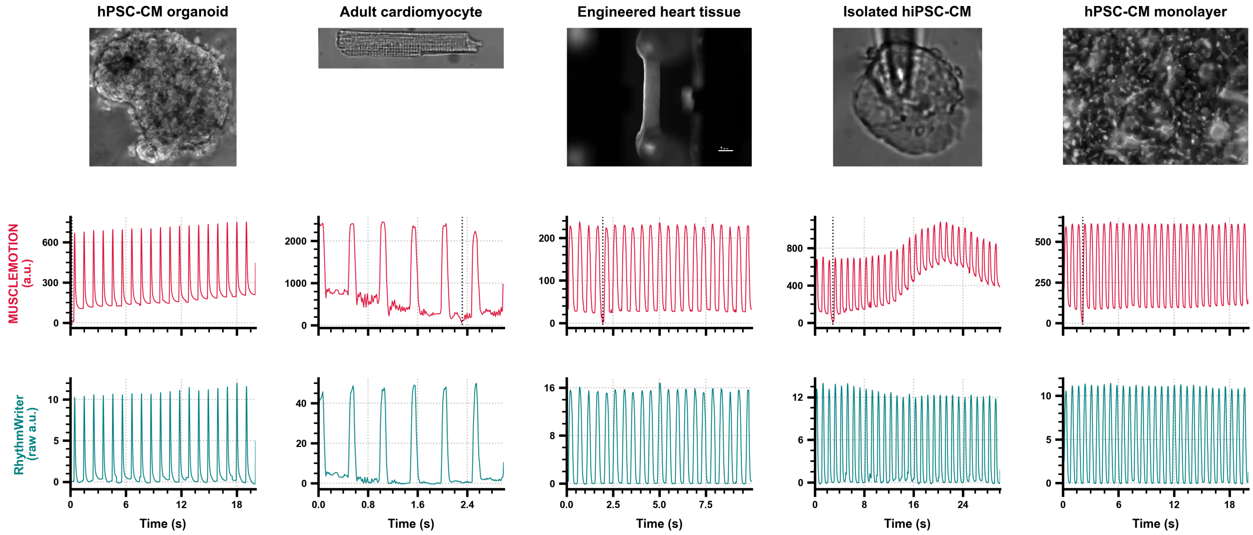


**Supplementary Figure S6| RhythmWriter is robust for multiple tissue types.** RhythmWriter and MUSCLEMOTION on five of the recordings published with MUSCLEMOTION [1], each analyzed whole field over its whole length. Top, the first frame of each recording. Middle, MUSCLEMOTION v1.0 with default settings (red). Dotted lines, its reference frame. Bottom, RhythmWriter (teal).

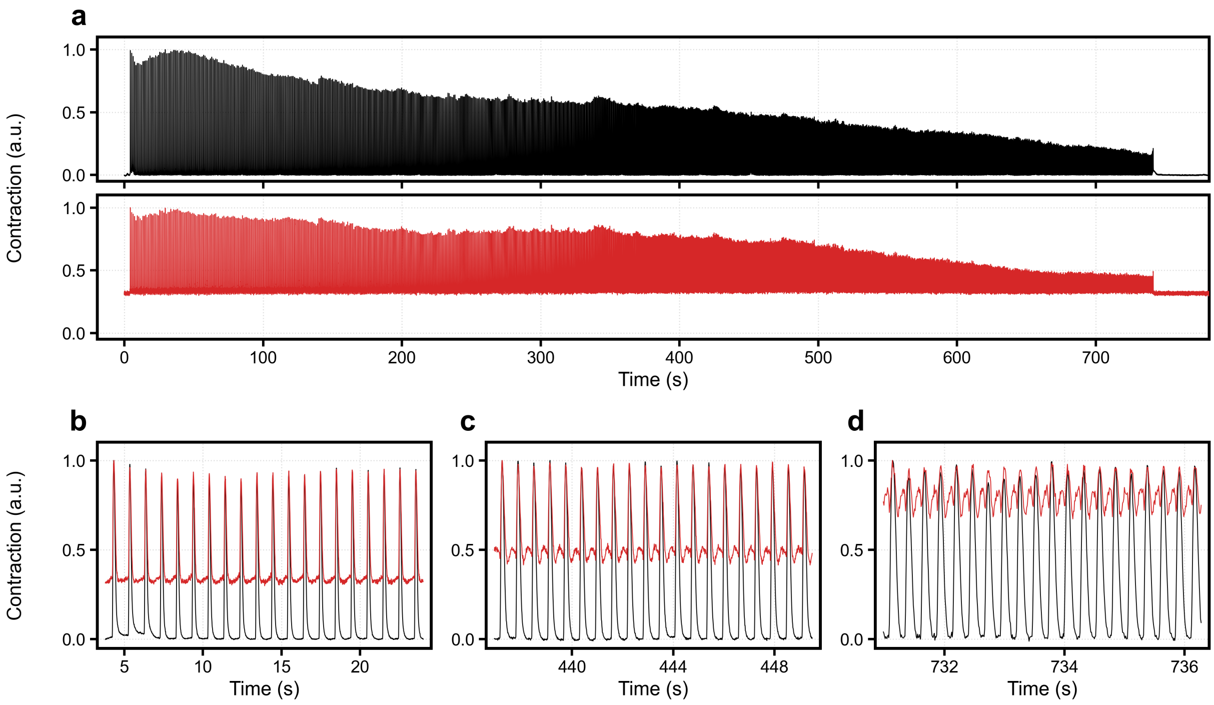


**Supplementary Figure S7| PLoRI is unstable when recovery intervals decrease.** Aggregate 2 (Fig. 5) under dynamic pacing, measured by RhythmWriter (black) and by PLoRI [2] v1.0.3 with default settings (red) on the same 134 × 134-pixel region. **a**, The whole recording (13.0 min), RhythmWriter above PLoRI, each scaled to its maximum. **b**–**d**, One step of 20 stimuli at programmed cycle lengths of 1000 ms (**b**), 618 ms (**c**) and 255 ms (**d**), each trace scaled to its maximum within the step. PLoRI's output at rest is 32% of its largest beat. As the beats shrink, PLoRI's median cycle minimum rises from 33% of its median cycle peak in **b** to 43% in **c** and 71% in **d**, and a second deflection grows in each cycle. RhythmWriter's stays within 1% of its peak at every step.
